# Heme oxygenase limits mycobacterial infection-induced ferroptosis

**DOI:** 10.1101/2021.05.04.442683

**Authors:** Kaiming Luo, Roland Stocker, Warwick J Britton, Kazu Kikuchi, Stefan H Oehlers

## Abstract

Iron homeostasis is essential for both sides of the host-pathogen interface. Restricting access of iron slows bacterial growth while iron is also a necessary co-factor for host immunity. Heme oxygenase 1 (HMOX1) is a critical regulator of iron homeostasis that catalyses the liberation of iron during degradation of heme. It is also a stress-responsive protein that can be rapidly upregulated and confers protection to the host. Although a protective role of HMOX1 has been demonstrated in a variety of diseases, the role of HMOX1 in *Mycobacterium tuberculosis* infection is equivocal across experiments with different host-pathogen combinations. Here we use the natural host-pathogen pairing of the zebrafish-*Mycobacterium marinum* infection platform to study the role of zebrafish heme oxygenase in mycobacterial infection. We identify zebrafish Hmox1a as the relevant functional paralog of mammalian HMOX1 and demonstrate a conserved role for Hmox1a in protecting the host from mycobacterial infection. Using genetic and chemical tools, we show zebrafish Hmox1a protects the host against mycobacterial infection by reducing infection-induced iron accumulation and ferroptosis.

## Introduction

Infection with pathogenic mycobacteria, such as *Mycobacterium tuberculosis* (*Mtb*), leads to the formation of granulomas, the hallmark histological feature of tuberculosis (TB) [1]. Host cells within granulomas undergo significant phenotypic remodelling including the upregulation of cytoprotective stress response proteins in response to mycobacterial virulence factors and changes to the microenvironment [2–4].

Haem oxygenase 1 (HMOX1), a key regulator of iron homeostasis and cellular redox biology, is expressed within human- and mouse-*Mtb* granulomas [3–9]. HMOX1-derived carbon monoxide inhibits the growth of mycobacteria by direct toxicity and inducing mycobacterial dormancy gene expression [10–12]. HMOX1-deficient mice are more susceptible to mycobacterial infection, while inhibition of HMOX1 activity with tin protoporphyrin (SnPP) increases hosts resistance to mycobacterial infection in human macrophages and mouse models [3–5, 8, 10, 13]. It is unclear if these contrasting effects on *Mtb* infection are a function of losing non-enzymatic HMOX1 functions in the gene deficient mice, differences in infection models, or other reasons [14].

HMOX1 is a highly evolutionarily conserved enzyme that has been identified in a wide variety of organisms. Zebrafish *hmox1a* and *hmox1b* encode paralogs of mammalian HMOX1, and their transcription is responsive to a range of oxidative stress-inducing conditions [15–17]. The zebrafish-*M. marinum* model has been widely used to study conserved host redox perturbations associated with mycobacterial infection [18–20]. Here we have used zebrafish to investigate the role of Hmox1a in the control of mycobacterial infection. We provide evidence that induction of host Hmox1a expression restricts iron supply to limit mycobacterial growth and prevent excessive ferroptosis.

## Materials and Methods

### Zebrafish husbandry

Adult zebrafish were maintained at Centenary Institute and embryos were obtained by natural spawning followed and raised in E3 media at 28-32°C (Sydney Local Health District AWC Approvals: 16-037 and 17-036).

### *M. marinum* infection

Single cell suspensions of mid log-phase fluorescent *M. marinum* strains and ΔESX1 *M. marinum* were stored at −80°C in aliquots [21]. Zebrafish infections were carried out as previously described with 200 CFU infection doses by microinjection into embryos or intraperitoneal injection into adults [21, 22]. Infected adults were maintained at 28°C with 14:10 light:dark lighting cycle.

### Quantitative Real–time PCR

Total RNA was extracted from homogenates using Trizol (Thermofisher) and cDNA was synthesised with a High Capacity cDNA Synthesis Kit (Applied Biosystems). qPCR reactions were performed on a LightCycler® 480 System. Gene expression was quantified by the delta–delta C_T_ method using normalisation to zebrafish *bact* or *M. marinum 18s* as appropriate. Sequences of primers are listed in Supplementary Table 1.

### Histology

Cryosectioning was performed as previously described [22]. Hmox1 immunostaining was carried out with a mouse anti-HMOX1 primary (GeneTex GTX633693) and a goat anti-mouse Alexa Fluor 488 (Thermofisher R37120). Perls’ Prussian Blue staining was performed in acid ferrocyanide solution (equal amount of 5% aqueous potassium ferrocyanide and 5% HCl) for 30 min at room temperature followed by two washes of distilled water and counterstaining with 0.1% nuclear fast red. Whole mount embryo Perls’ Prussian Blue staining was performed without the counterstaining step.

### CRISPR/Cas9 Gene Editing Technique

The gRNA target sites for each gene were designed using CRISPRscan. The sequences of gRNA oligonucleotides are listed in Supplementary Table 2. The gRNA templates were amplified by PCR with scaffold reverse primer, and then transcribed with HiScribe™ T7 High Yield RNA Synthesis Kit (NEB) [23]. One cell stage embryos were injected with 1 nL of a mixture containing 200 ng/μL of the four gRNAs and 2 ng/μL Cas9.

### Imaging

Imaging was carried out on Leica M205FA and DM6000B, and Deltavision Elite microscopes as previously described [18, 21, 22].

### Image analysis

Fluorescent pixel count for enumeration of bacterial burden and quantification of fluorescent staining was carried out as previously described in ImageJ [21]. Fluorescent stain areas are reported as pixels per granuloma.

Perls’ Prussian Blue staining in embryos was quantified by splitting the blue channel from colour images and then using the “Measure” function in ImageJ to quantify the inverse mean pixel intensity of a constant area within granulomas. The “average blue intensity per granuloma” was calculated by subtracting the background blue pixel intensity from the granuloma blue pixel intensity.

### Whole mount in situ hybridisation

Sequences of primers used to generate the DIG-labelled *hmox1a* probe are listed in Supplementary Table 1, *in situ* hybridisation was carried out as previously described [24].

### Cell death and reactive oxygen species staining

TUNEL and CellROX staining were performed as previously described and according to manufacturer’s instructions [18].

### Statistical analysis

Student’s *t* and ANOVA tests were carried out as appropriate for multiple comparisons using Graphpad Prism. Each data point indicates a single animal unless otherwise stated. Data are plotted as means +/− standard deviation.

## Results

### Zebrafish *hmox1a* is the functional Hmox paralog in *M. marinum* infection

Zebrafish have four Hmox-encoding paralogs: *hmox1a*, *hmox1b*, *hmox2a*, and *hmox2b* [15–17]. To determine which paralog is functional in *M. marinum* infection, we first infected adult zebrafish by intraperitoneal injection and analysed gene expression changes at 14 days post infection (dpi). Infected adults had increased expression of *hmox1a*, but not the other paralogs (Fig. 1A).

**Figure 1:**
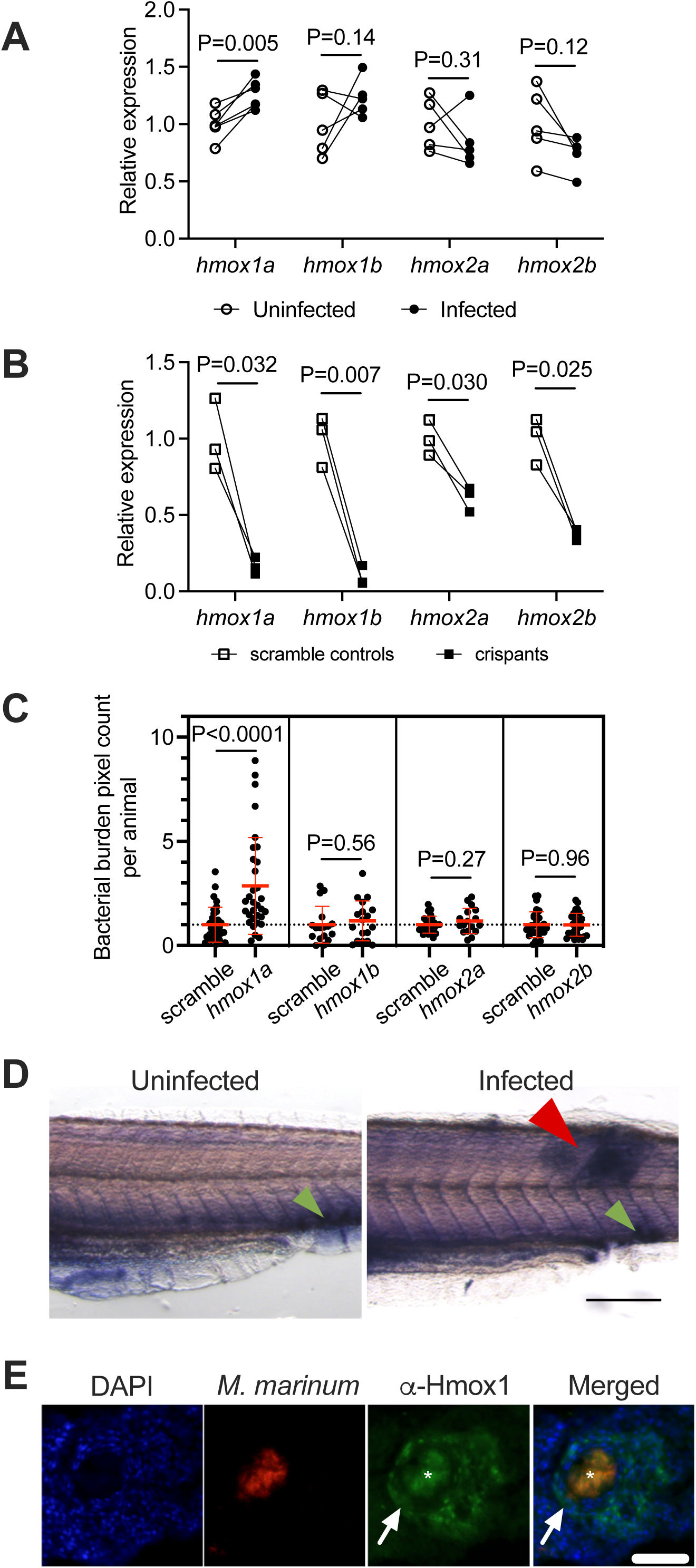
Zebrafish *hmox1a* is the functional paralog of mammalian *HMOX1* during mycobacterial infection A. qPCR expression analysis of zebrafish *hmox* paralogs in homogenates of 14 dpi adult zebrafish infected with *M. marinum*. Biological pairs indicated by matching lines. B. qPCR expression analysis of zebrafish *hmox* paralogs in 7 dpf crispants. Biological pairs indicated by matching lines. C. *M. marinum* burden measured by fluorescent pixel count in 5 dpi crispants. D. Whole mount *in situ* hybridisation detection of *hmox1a* transcripts in 5 dpi zebrafish embryos. Green arrowhead indicates site of constitutive CHT expression, red arrowhead indicates site of inducible expression at a granuloma. Scale bar represents 200 μm. E. Representative image of antibody detection of Hmox1 in a granuloma of a 14 dpi adult zebrafish. Arrows indicate cellular rim of granuloma. Asterisk indicates bleed through fluorescence of *M. marinum*-TdTomato signal in necrotic core. Scale bar represents 100 μm.

To investigate the function of each paralog during *M. marinum* infection, we next used CRISPR/Cas9 technology to knockdown each paralog and infected crispant embryos with *M. marinum* (Fig. 1B). Knockdown of *hmox1a* increased the bacterial burden at 5 dpi, while knockdown of *hmox1b*, *hmox2a*, and *hmox2b* did not affect bacterial burden (Fig. 1C).

To investigate *hmox1a* expression in more detail, whole mount *in situ* hybridisation was used to visualise the spatial distribution of *hmox1a* expression within *M. marinum-*infected embryos. *hmox1a* was highly expressed in the haematopoietic niche of the caudal haematopoietic tissue and around granulomas (Fig. 1D). To spatially examine Hmox expression in adult granulomas, we used a mouse Hmox1 antibody which would potentially detect both Hmox1a and Hmox1b in zebrafish. Hmox1 staining was detected within the host cellular rim of granulomas from 14 dpi zebrafish adults (Fig. 1E).

### Hmox1a-dependent *M. marinum* granuloma formation does not explain increased susceptibility to infection

To investigate the function of *hmox1a* in the formation of adult zebrafish-*M. marinum* granulomas, we utilised our *hmox1a^vcc42^* knockout allele [25]. As expected from our CRISPR-Cas9 knockdown studies, *hmox1a^vcc42/vcc42^* embryos displayed significantly increased bacterial burden compared to WT clutch mates (Fig. 2A). Adult *hmox1a^vcc42/vcc42^* mutants had increased mortality following infection that was consistent with the increased bacterial burden (Fig. 2B). Heterozygous *hmox1a^+/vcc42^* zebrafish displayed intermediate phenotypes in both assays (Fig. 2A and B).

**Figure 2:**
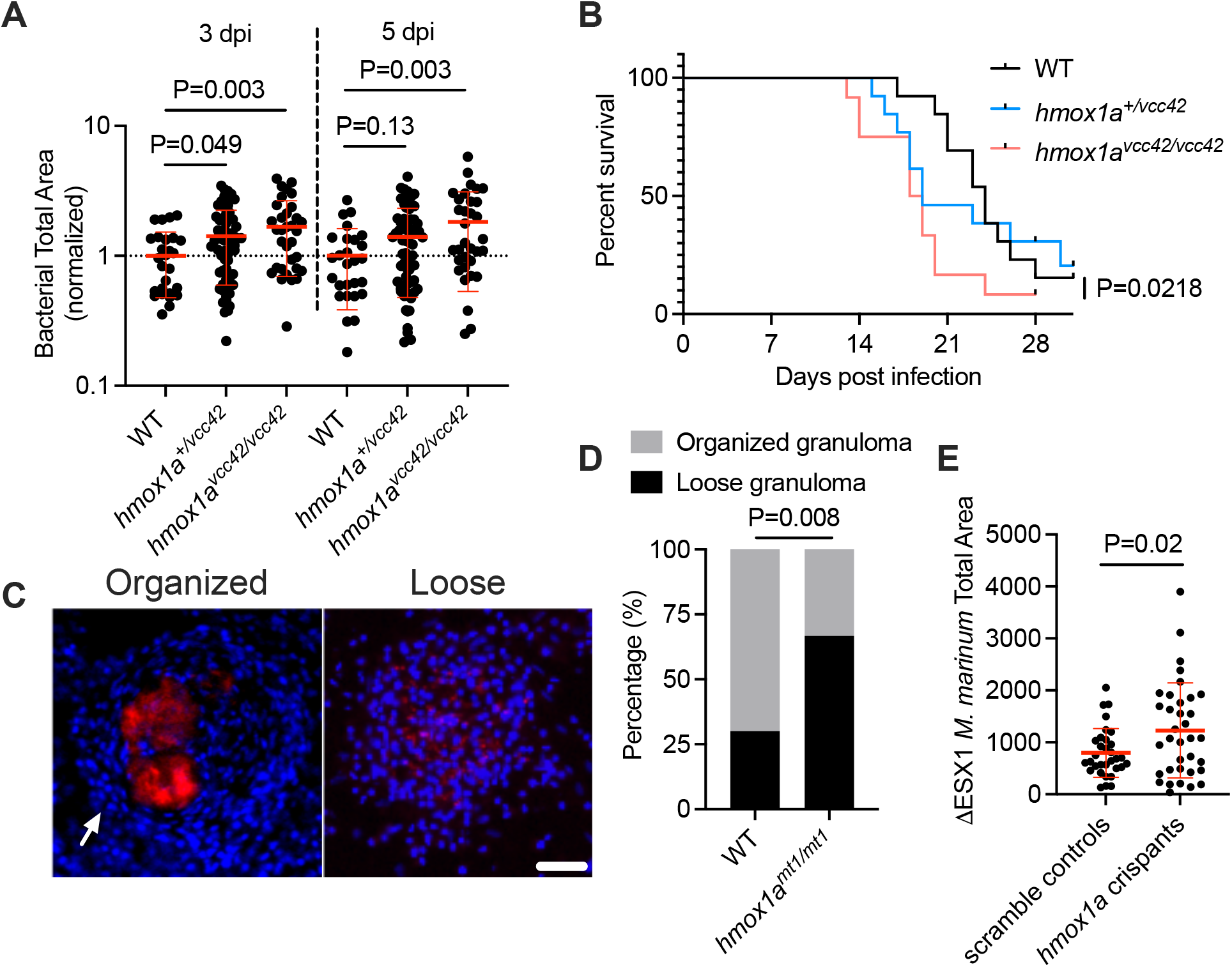
Zebrafish Hmox1a is necessary for efficient granuloma formation A. Bacterial burden in 5 dpi *hmox1a^vcc41/vcc42^* mutant embryos. B. Survival of adult *hmox1a^vcc41/vcc42^* mutant zebrafish following infection with *M. marinum*. n=13 WT, 13 heterozygous mutants, 12 homozygous mutants. C. Representative images of granuloma morphology classes in 14 dpi *hmox1a^vcc41/vcc42^* mutant adult zebrafish. Arrow indicates organised cellular rim of granuloma. Scale bar represents 50 μm. D. Quantification of granuloma morphology classes in 14 dpi *hmox1a^vcc41/vcc42^* mutant adult zebrafish. n = 40 WT granulomas from 3 animals, 24 *hmox1a^vcc41/vcc42^* mutant granulomas from 3 animals. E. Bacterial burden in *hmox1a* crispants infected with ΔESX1 *M. marinum*.

Failure to form granulomas has been proposed to underly the mycobacterial control defect in *Hmox1*-null mice [3, 4]. To determine if zebrafish reproduced this phenotype, we scored *M. marinum* granulomas as “organised” or “loose” based on host nuclear structure in *hmox1a^vcc42/vcc42^* and their WT clutch mates (Fig. 2C) [2, 22]. The proportion of unorganised “loose” granulomas in *hmox1a^vcc42/vcc42^* adults was significantly higher than in the WT adults demonstrating a conserved defect in granuloma formation (Fig. 2D).

To investigate the association between *hmox1a* and granuloma formation, we infected *hmox1a^vcc42/vcc42^* embryos with ΔESX1 *M. marinum*, a strain that is unable to drive granuloma formation [26]. Unexpectedly, ΔESX1 *M. marinum* load was significantly increased in *hmox1a^vcc42/vcc42^* homozygous larvae at both 3 and 5 dpi suggesting a more general susceptibility to mycobacterial infection than a defect in granuloma formation alone (Fig. 2E).

### Hmox1a restricts iron to control mycobacterial infection

Since HMOX1 is a key regulator of iron homeostasis, we performed Perls’ Prussian blue staining on sections from WT and *hmox1a^vcc42/vcc42^* adults to visualise free iron (Fig. 3A). There were more Perls’ Prussian blue positive granulomas in *hmox1a^vcc42/vcc42^* mutants than WT zebrafish (Fig. 3B). Perls’ Prussian blue staining of *M. marinum*-infected embryos was not as vivid as that of adult granulomas (Fig. 3C), but spectral quantification staining detected more Perls’ Prussian blue staining in granulomas from *hmox1a* crispants (Fig. 3D).

**Figure 3:**
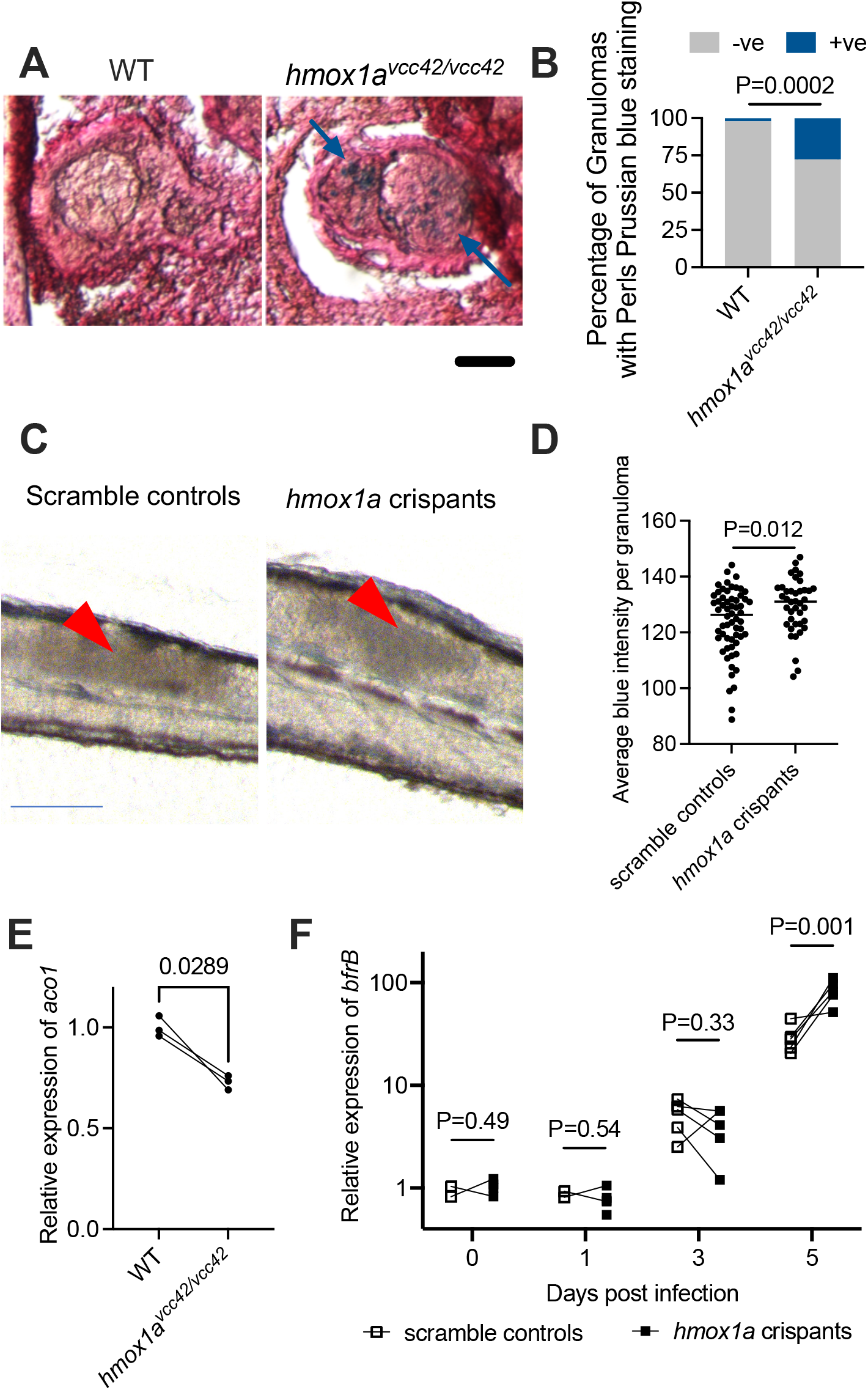
Zebrafish Hmox1a restricts iron availability during infection A. Representative images of Perls’ Prussian blue staining of granulomas from 14 dpi *hmox1a^vcc41/vcc42^* mutant adult zebrafish. Blue arrows indicate location of positive staining. Scale bar represents 50 μm. B. Quantification of granuloma Perls’ Prussian blue staining in *hmox1a^vcc41/vcc42^* mutant adult zebrafish. n = 46 granulomas from 3 WT animals, 83 granulomas from 3 *hmox1a^vcc41/vcc42^* mutants. C. Representative images of Perls’ Prussian blue staining of 5 dpi zebrafish embryos. Red arrows indicate locations of analysed granulomas. Scale bar represents 100 μm. D. Quantification of granuloma Perls’ Prussian blue staining in 5 dpi *hmox1a* crispants. Each data point represents a single granuloma from an individual embryo. E. qPCR analysis of host *aco1* gene expression in 5 dpi *hmox1a^vcc41/vcc42^* mutant embryos. F. qPCR analysis of *M. marinum bfrB* gene expression in *hmox1a* crispants.

To confirm the presence of increased iron at the host-pathogen interface, we performed gene expression analysis of *aco1*, a host iron-suppressed regulator of iron metabolism [27], which was downregulated in 5 dpi *hmox1a^vcc42/vcc42^* embryos (Fig. 3E). To investigate the bacterial response, we profiled the expression of *bfrB*, an iron storage gene in *M. marinum* homologous to human ferritin. Expression of *bfrB* was progressively increased from 1 to 3 to 5 dpi and was higher in *hmox1a* crispants than WT hosts at 5 dpi (Fig. 3F).

### Hmox1a-deficiency exposes hosts to infection-induced ferroptosis

Mycobacteria require iron as a redox cofactor for vital enzymes and utilise multiple strategies to acquire iron from the host [28, 29]. Excessive iron can be toxic to host cells by triggering ferroptosis, a recently described mode of cell death associated with lipid peroxidation that drives pathology in the mouse-*Mtb* model [30, 31]. We hypothesised ferroptosis could be responsible for the increased *M. marinum* burden in our Hmox1a-deficent zebrafish.

Consistent with the increased iron in Hmox1a-deficient zebrafish, we observed increased granuloma CellROX staining in *hmox1a* crispants compared to control embryos (Fig. 4A). We also observed more TUNEL positive cells around granulomas in *hmox1a* crispants (Fig. 4B).

**Figure 4:**
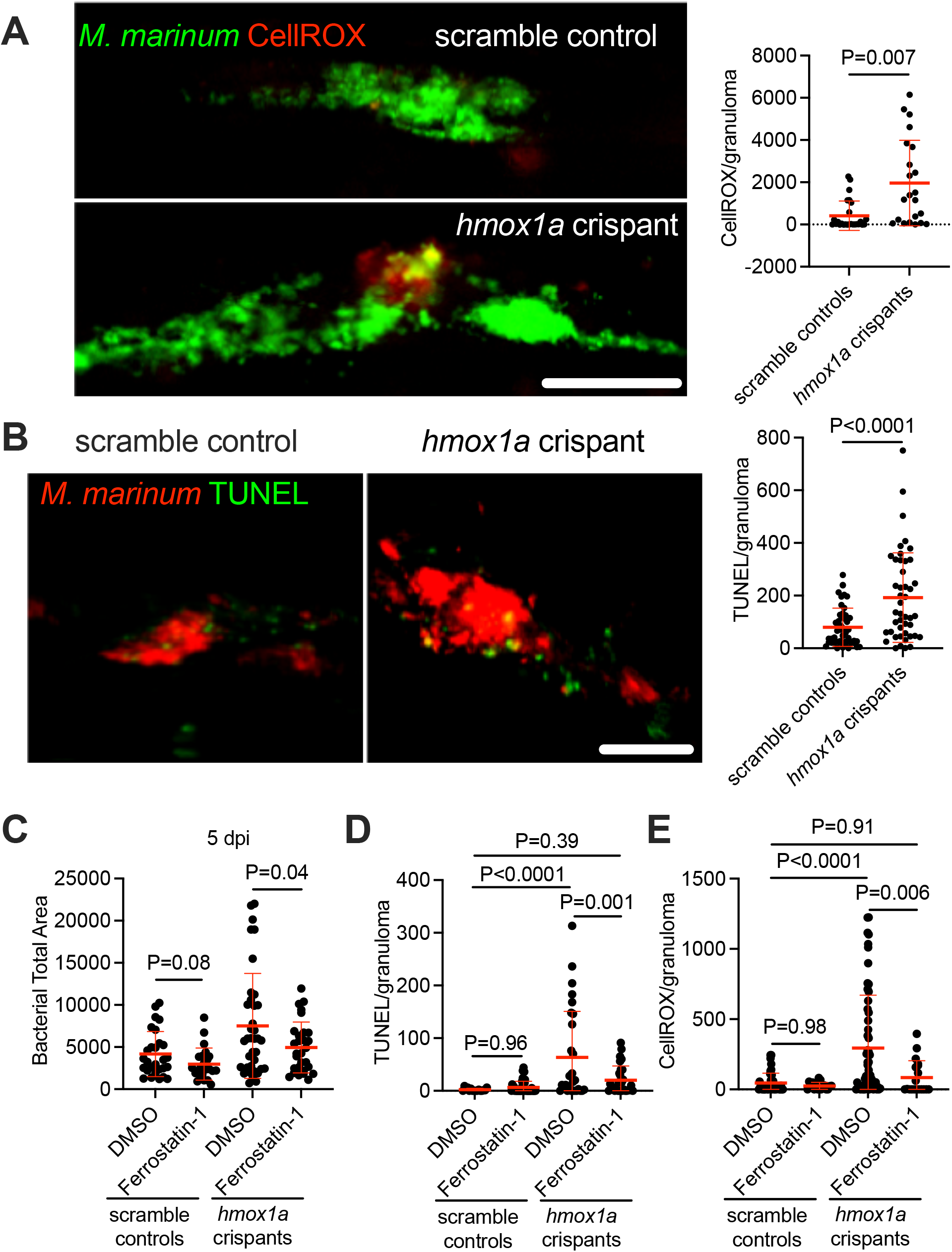
Zebrafish Hmox1a prevents infection-induced ferroptosis A. Representative images and quantification of CellROX staining in 5 dpi *hmox1a* crispants. Scale bar represents 50 μm. B. Representative images and quantification of TUNEL staining in 5 dpi *hmox1a* crispants. Scale bar represents 50 μm. C. Bacterial burden in 5 dpi *hmox1a* crispants treated with ferrostatin-1. D. Quantification of CellROX staining in 5 dpi *hmox1a* crispants treated with ferrostatin-1. E. Quantification of TUNEL staining in 5 dpi *hmox1a* crispants treated with ferrostatin-1.

To test the role of ferroptosis in these phenotypes we treated embryos with ferrostatin, a small molecule inhibitor of ferroptosis. Ferrostatin treatment reduced bacterial burden in *hmox1a* crispants but not in scramble controls (Fig. 4C). Ferrostatin treatment also reduced CellROX staining in granulomas and the number of TUNEL positive cells per granuloma (Fig 4D and 4E).

## Discussion

Host cells need to balance iron availability and the production of reactive oxygen species during infection with intracellular pathogens. Our data show infection-induced Hmox1a is a host-protective enzyme that aids containment of infection within granulomas and slows the growth of mycobacteria by restricting iron availability and preventing ferroptosis.

Our study clearly identifies zebrafish *hmox1a* as the functional ortholog of mammalian HMOX1 as the most transcriptionally responsive to infection in adult zebrafish and the only gene which had a knockdown burden phenotype in the CRIPSR-Cas9 embryo infection model. Our data further suggests zebrafish *hmox1b* does not functionally compensate for loss of *hmox1a* and may be a pseudogene.

Our experiment infecting *hmox1a*-deficient zebrafish with the ΔESX1 *M. marinum* suggests *hmox1a* is involved in control of mycobacterial infection upstream of granuloma formation. We found iron accumulation in mycobacterial granulomas which correlated with CellROX staining and increased cell death in *hmox1a*-deficient zebrafish. These phenotypes were reversed after treatment with ferrostatin suggesting the mycobacterial control defect in *hmox1a*-deficient zebrafish could be driven by iron-induced ferroptosis upstream of granuloma formation.

The non-specific action of ferrostatin-1 is a potential limitation of our study. Ferrostatin-1 has been recently reported to inhibit the enzymic activity of 15-Lipoxygenase, which produces the HpETE-PE ferroptotic cell death signal as part of the 15LOX/PEBP1 complex with iron as a cofactor [32, 33]. Infection-induced lipoxygenase and cyclooxygenase activity in mycobacterial infection generally leads to unfavourable suppression of inflammation, while inhibition of these enzyme families generally controls mycobacterial infection [34–38]. Thus, we are unable to rule out 15LOX/PEBP1-induced ferroptosis as a possible mechanism of ferrostatin-induced protection in our *hmox1a*-deficient zebrafish.

The lack of host protection in WT embryos treated with ferrostatin suggests ferroptosis is of limited importance in the zebrafish embryo-*M. marinum* infection model. Our data show that depletion of Hmox1a is required to sensitise embryos to biologically significant levels of ferroptosis by impairing endogenous iron metabolism.

The results from the infection time course highlights 3-5 dpi as a key window in *M. marinum* infection when granulomas undergo extensive organisation and expansion. During this time period, host *hmox1a* and bacterial *bfrB* are highly upregulated, and *hmox1a*-deficient zebrafish embryos develop their conditional pathologies. Together, these data provide evidence of heme oxygenase function in the race to acquire iron between host and mycobacterium during granuloma formation.

## Acknowledgements

We thank Drs Elinor Hortle and Pradeep Cholan for zebrafish infection and analysis methods training, Drs Angela Kurz and Kristina Jahn of Sydney Cytometry for assistance with imaging equipment, the Victor Chang Cardiac Research Institute BioCORE staff for zebrafish maintenance, members of the Tuberculosis Research Program at the Centenary Institute for helpful discussion.

## Funding

This work was supported by a Chinese Scholarships Council Postgraduate Scholarship to KL; the University of Sydney Fellowship G197581, NSW Ministry of Health under the NSW Health Early-Mid Career Fellowships Scheme H18/31086 to SHO; the NHMRC Centre of Research Excellence in Tuberculosis Control (APP1153493) to WJB.

## Notes

### Competing Interest Statement

The authors have declared no competing interest.

### Summary of Updates

updated references

## References

1. Ramakrishnan, L., Mycobacterium tuberculosis pathogenicity viewed through the lens of molecular Koch’s postulates. Curr Opin Microbiol, 2020. 54: p. 103–110.

2. Cronan, M.R., et al., Macrophage Epithelial Reprogramming Underlies Mycobacterial Granuloma Formation and Promotes Infection. Immunity, 2016. 45(4): p. 861–876.

3. Regev, D., et al., Heme oxygenase-1 promotes granuloma development and protects against dissemination of mycobacteria. Lab Invest, 2012. 92(11): p. 1541–52.

4. Silva-Gomes, S., et al., Heme catabolism by heme oxygenase-1 confers host resistance to Mycobacterium infection. Infect Immun, 2013. 81(7): p. 2536–45.

5. Costa, D.L., et al., Pharmacological Inhibition of Host Heme Oxygenase-1 Suppresses Mycobacterium tuberculosis Infection In Vivo by a Mechanism Dependent on T Lymphocytes. MBio, 2016. 7(5).

6. Abdalla, M.Y., et al., Induction of heme oxygenase-1 contributes to survival of Mycobacterium abscessus in human macrophages-like THP-1 cells. Redox Biol, 2015. 4: p. 328–39.

7. Rockwood, N., et al., Mycobacterium tuberculosis Induction of Heme Oxygenase-1 Expression Is Dependent on Oxidative Stress and Reflects Treatment Outcomes. Front Immunol, 2017. 8: p. 542.

8. Chinta, K.C., et al., Microanatomic Distribution of Myeloid Heme Oxygenase-1 Protects against Free Radical-Mediated Immunopathology in Human Tuberculosis. Cell Rep, 2018. 25(7): p. 1938–1952 e5.

9. Scharn, C.R., et al., Heme Oxygenase-1 Regulates Inflammation and Mycobacterial Survival in Human Macrophages during Mycobacterium tuberculosis Infection. J Immunol, 2016. 196(11): p. 4641-9.

10. Shiloh, M.U., P. Manzanillo, and J.S. Cox, Mycobacterium tuberculosis senses host-derived carbon monoxide during macrophage infection. Cell Host Microbe, 2008. 3(5): p. 323–30.

11. Kumar, A., et al., Heme oxygenase-1-derived carbon monoxide induces the Mycobacterium tuberculosis dormancy regulon. J Biol Chem, 2008. 283(26): p. 18032–9.

12. Zacharia, V.M., et al., cor, a novel carbon monoxide resistance gene, is essential for Mycobacterium tuberculosis pathogenesis. mBio, 2013. 4(6): p. e00721–13.

13. Costa, D.L., et al., Heme oxygenase-1 inhibition promotes IFNgamma- and NOS2-mediated control of Mycobacterium tuberculosis infection. Mucosal Immunol, 2021. 14(1): p. 253–266.

14. Costa, D.L., et al., Modulation of Inflammation and Immune Responses by Heme Oxygenase-1: Implications for Infection with Intracellular Pathogens. Antioxidants (Basel), 2020. 9(12).

15. Mills, M.G. and E.P. Gallagher, A targeted gene expression platform allows for rapid analysis of chemical-induced antioxidant mRNA expression in zebrafish larvae. PLoS One, 2017. 12(2): p. e0171025.

16. Holowiecki, A., B. O’Shields, and M.J. Jenny, Spatiotemporal expression and transcriptional regulation of heme oxygenase and biliverdin reductase genes in zebrafish (Danio rerio) suggest novel roles during early developmental periods of heightened oxidative stress. Comp Biochem Physiol C Toxicol Pharmacol, 2017. 191: p. 138–151.

17. Holowiecki, A., B. O’Shields, and M.J. Jenny, Characterization of heme oxygenase and biliverdin reductase gene expression in zebrafish (Danio rerio): Basal expression and response to pro-oxidant exposures. Toxicol Appl Pharmacol, 2016. 311: p. 74–87.

18. Black, H.D., et al., The cyclic nitroxide antioxidant 4-methoxy-TEMPO decreases mycobacterial burden in vivo through host and bacterial targets. Free Radic Biol Med, 2019. 135: p. 157–166.

19. Roca, F.J. and L. Ramakrishnan, TNF Dually Mediates Resistance and Susceptibility to Mycobacteria via Mitochondrial Reactive Oxygen Species. Cell, 2013. 153(3): p. 521–34.

20. Roca, F.J., et al., TNF Induces Pathogenic Programmed Macrophage Necrosis in Tuberculosis through a Mitochondrial-Lysosomal-Endoplasmic Reticulum Circuit. Cell, 2019. 178(6): p. 1344–1361 e11.

21. Matty, M.A., S.H. Oehlers, and D.M. Tobin, Live Imaging of Host-Pathogen Interactions in Zebrafish Larvae. Methods Mol Biol, 2016. 1451: p. 207–23.

22. Cheng, T., et al., High content analysis of granuloma histology and neutrophilic inflammation in adult zebrafish infected with Mycobacterium marinum. Micron, 2020. 129: p. 102782.

23. Wu, R.S., et al., A Rapid Method for Directed Gene Knockout for Screening in G0 Zebrafish. Dev Cell, 2018. 46(1): p. 112–125 e4.

24. Thisse, C. and B. Thisse, High-resolution in situ hybridization to whole-mount zebrafish embryos. Nat Protoc, 2008. 3(1): p. 59–69.

25. Luo, K., et al., Zebrafish heme oxygenase 1a is necessary for normal development and macrophage migration. bioRxiv, 2021: p. 2021.04.07.438802.

26. Volkman, H.E., et al., Tuberculous granuloma formation is enhanced by a mycobacterium virulence determinant. PLoS Biol, 2004. 2(11): p. e367.

27. Pantopoulos, K., Iron metabolism and the IRE/IRP regulatory system: an update. Ann N Y Acad Sci, 2004. 1012: p. 1–13.

28. Marcela Rodriguez, G. and O. Neyrolles, Metallobiology of Tuberculosis. Microbiol Spectr, 2014. 2(3).

29. Pandey, R. and G.M. Rodriguez, A ferritin mutant of Mycobacterium tuberculosis is highly susceptible to killing by antibiotics and is unable to establish a chronic infection in mice. Infect Immun, 2012. 80(10): p. 3650–9.

30. Amaral, E.P., et al., A major role for ferroptosis in Mycobacterium tuberculosis-induced cell death and tissue necrosis. J Exp Med, 2019. 216(3): p. 556–570.

31. Dixon, S.J., et al., Ferroptosis: an iron-dependent form of nonapoptotic cell death. Cell, 2012. 149(5): p. 1060–72.

32. Anthonymuthu, T.S., et al., Resolving the paradox of ferroptotic cell death: Ferrostatin-1 binds to 15LOX/PEBP1 complex, suppresses generation of peroxidized ETE-PE, and protects against ferroptosis. Redox Biol, 2021. 38: p. 101744.

33. Kagan, V.E., et al., Oxidized arachidonic and adrenic PEs navigate cells to ferroptosis. Nat Chem Biol, 2017. 13(1): p. 81–90.

34. Hortle, E., et al., Thrombocyte Inhibition Restores Protective Immunity to Mycobacterial Infection in Zebrafish. J Infect Dis, 2019. 220(3): p. 524–534.

35. Bafica, A., et al., Host control of Mycobacterium tuberculosis is regulated by 5-lipoxygenase-dependent lipoxin production. J Clin Invest, 2005. 115(6): p. 1601–6.

36. Chen, M., et al., Lipid mediators in innate immunity against tuberculosis: opposing roles of PGE2 and LXA4 in the induction of macrophage death. J Exp Med, 2008. 205(12): p. 2791–801.

37. Lewis, A. and P.M. Elks, Hypoxia Induces Macrophage tnfa Expression via Cyclooxygenase and Prostaglandin E2 in vivo. Front Immunol, 2019. 10: p. 2321.

38. Tobin, D.M., et al., Host genotype-specific therapies can optimize the inflammatory response to mycobacterial infections. Cell, 2012. 148(3): p. 434–46.

